# Sensory capability and information integration independently predict cognitive status of healthy older adults

**DOI:** 10.1101/2020.04.20.050344

**Authors:** Jonas Misselhorn, Florian Göschl, Focko L. Higgen, Friedhelm C. Hummel, Christian Gerloff, Andreas K. Engel

## Abstract

Ageing is characterized by changes in sensory and cognitive abilities. While there is evidence that decline in sensory acuity and enhanced multisensory integration predict cognitive status in healthy older adults, potential mechanistic links between these age-related alterations remain unclear. In the current study, we assessed performance of younger and older healthy adults in a visuotactile delayed match-to-sample task and related indices of multisensory integration to unisensory perceptual thresholds and cognitive assessment data. Additionally, we applied transcranial alternating current stimulation (tACS) to modulate cortical networks found to underlie visuotactile interactions and working-memory matching in our previous work. Analysing response times and signal detection measures, we found older adults to show enhanced multisensory integration and benefit more from successful working memory matching. Both measures predicted cognitive status and correlated positively with each other, suggesting that they likely reflect a common underlying tendency to integrate information. Sensory capability, however, independently predicted cognitive status. tACS with beta frequency (20 Hz) accelerated task performance and this effect was more pronounced in the older group. We conclude that sensory capability and information integration represent independent predictors of cognitive status. Finally, we discuss a potential role of the parietal cortex in mediating augmented integration in older adults.

## Introduction

Ageing is associated with marked alterations in sensory and cognitive abilities that are linked to changes in the integrity of the peripheral and central nervous system. Most detrimental to independence in daily live is a decline in ‘higher-level’ cognitive functions such as attention or working memory^1, 2^. While a number of theories on cognitive ageing focus on changes in frontal cortices^3–5^, an intriguing hypothesis proposes that decline in sensory capabilities – resulting from changes in peripheral sensory organs and sensory cortices – might mediate cognitive decline^6^. This idea is based on the established finding that simple measures of sensory acuity reliably predict cognitive abilities in older adults^6–8^.

Another well-documented observation is that older adults show enhanced multisensory integration, potentially to compensate for a decline in sensory acuity^9–12^. Furthermore, it has been shown that enhanced integration between modalities and sensory dominance in multisensory contexts jointly predict cognitive status of older adults^13^. It is an open question how sensory capability and multisensory integration interact as a function of age, and whether sensory decline might evoke enhanced integration in order to compensate for the loss in sensory acuity. Specifically, it is possible that these two predictors of cognitive abilities are linked and thus would explain the same variability in age-related cognitive decline. Alternatively, sensory capabilities and the tendency to integrate across the senses might be independent and thus point towards distinct processes of brain ageing.

In the current study, we employed a variant of a well-established visuotactile task^14,15^ to investigate multisensory integration in healthy younger and older adults. We assessed the relation of multisensory processes and (uni-)sensory capability as well as their potential to predict cognitive status. Participants identified tactile dot patterns, similar to Braille letters, and matched them with a target pattern held in working memory (Figure 1 A, B). In one third of the trials, tactile stimuli were presented unimodally, whereas two thirds featured additional task-irrelevant visual input that could either be congruent or incongruent to the tactile stimulus. Building on previous findings^9,16,17^, we hypothesized that older compared to younger adults show enhanced multisensory integration and, therefore, profit more from redundant but congruent visual input in tactile target detection. Preceding the experiment, participants underwent neurological examination including elaborate psychometric thresholding as well as cognitive assessment allowing to investigate potential dependencies between multisensory integration, sensory capability and cognitive status.

**Figure 1.**
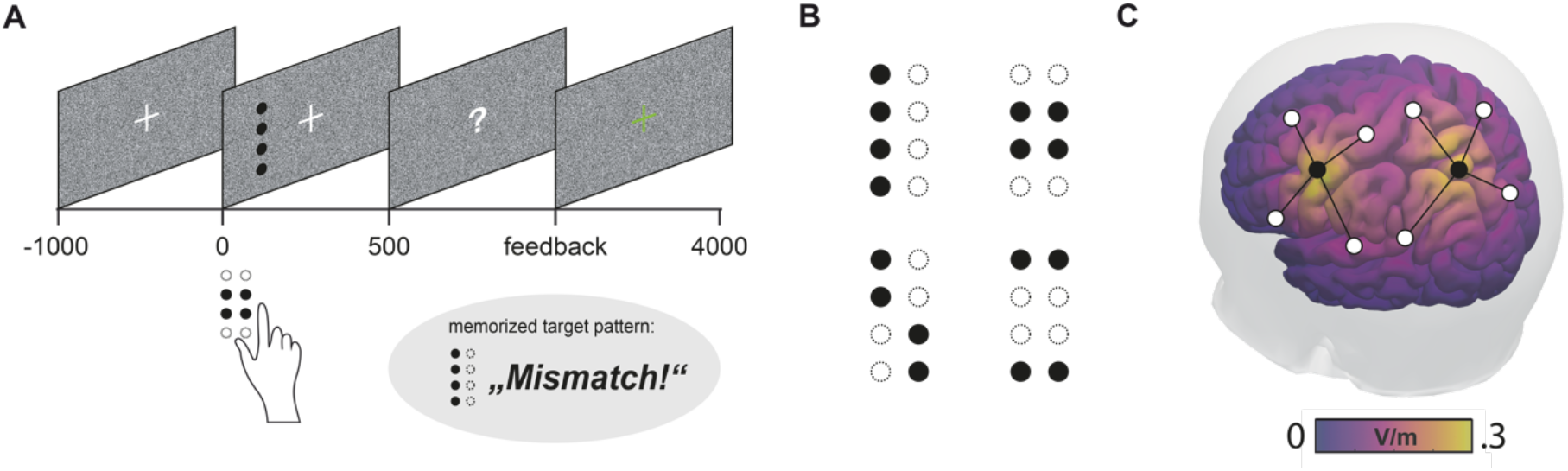
Experimental design. **(A)** Trial sequence. Participants fixated a central cross. In two thirds of all trials, tactile pattern presentation to the right index finger was accompanied by a visual dot pattern appearing on the left of the fixation cross. Participants were instructed to attend to the tactile input only, compare the tactile pattern with a blockwise-defined target pattern and report match or mismatch. **(B)** Set of dot patterns that were presented on a Braille stimulator and as a visual pattern on the screen. **(C)** tACS was applied using Ag/AgCl ring electrodes in two separate montages, centered on S1 (primary somatosensory cortex) and IPS (intraparietal sulcus) of the left hemisphere.

In addition, we used bifocal transcranial alternating current stimulation (tACS) in order to modulate corticocortical interactions. TACS has been rarely used to study older populations^18^ but has for instance been shown to improve age-related deficits in working memory^19^. Previous work suggested that the integration of information in multisensory brain networks might be realized via oscillatory activity^20, 21^ which in turn has been shown to profoundly change with age^22–24^. In a recent electroencephalography (EEG) study investigating visuotactile pattern matching, we had been able to describe a left-hemispheric cortical network mediating multisensory interactions and working memory that synchronised in the beta frequency range (around 20 Hz)^25^. Based on these findings, we stimulated a left-hemispheric network comprising primary somatosensory cortex (S1) and intraparietal sulcus (IPS; see Figure 1C) in the current study. We expected that beta-frequency stimulation (20 Hz) would enhance stimulus processing and working memory matching and lead to improved detection performance. We controlled tACS effects of beta stimulation with a sham condition as well as with an active control condition (70 Hz).

## Results

### Assessment

Preceding the experiment, participants underwent an assessment procedure, consisting of a neurological examination, the 2-point-discrimination test^26^ to assess peripheral somatosensation and a test of visual acuity (Snellen chart)^27^. Cognitive status was assessed with the Mini-Mental State Examination (MMSE)^28^ and the DemTect^29^. Using a visual analogue scale (VAS), participants further self-assessed attention level and fatigue. In the following, we report results of this assessment complemented by tactile and visual perception thresholds for the stimuli used in our task. These were estimated in a preceding study^15^. The group comparison of the assessment data (Table 1) showed significant differences between younger and older participants (Pillai’s Trace = 0.90, F = 22.47, *p* < 0.01). Post-hoc comparisons showed that the two groups differed significantly in tactile (*p* < 0.01) and visual (*p* < 0.01) thresholds. Furthermore, there was a significant difference in visual acuity (*p* < 0.01), but no difference in 2-point discrimination (*p* = 1.0). Despite higher variance in the older group, there were no differences in MMSE (*p* = 1.0) or DemTect (p = 0.23).

**Table 1.** Assessment data of the groups. Mean values are shown ± standard deviation. Based on significant main effects, post-hoc tests were conducted. ** indicate significant differences between younger and older participants, all significant *p*-values ≤ 0.01

| Metric | Younger group (n=17) | Older group (n=16) |
| --- | --- | --- |
| Age (years) | 24.4 (± 3.1) | 72.6 (± 4.6) |
| Gender | 10 females (7 males) | 10 females (6 males) |
| Tactile threshold (pin height in mm) | 0.55 (± 0.18)** | 1.21 (± 0.29)** |
| Visual threshold (grey intensity in RGB) | 48.1 (± 1.3)** | 55.8 (± 2.3)** |
| Visual acuity (arcmin) | 0.97 (± 0.1)** | 0.68 (± 0.1)** |
| 2-point-discrimination (mm) | 2.06 (± 0.2) | 2.13 (± 0.3) |
| MMSE | 29.7 (± 0.5) | 29.5 (± 1.0) |
| DemTect | 17.4 (± 1.5) | 16.3 (± 1.7) |
| Attention level (VAS in cm) | 1.5 (± 1.2) | 2.0 (± 1.6) |
| Fatigue (VAS in cm) | 1.8 (± 1.3) | 1.9 (± 1.7) |
| Fatigue of hand (VAS in cm) | 0.6 (± 0.6) | 1.3 (± 1.3) |

### Behaviour

Both detection accuracies and response times were analysed according to a mixed repeated measures ANOVA design with the following factors: AGE (between-participants, younger/older), STIMULUS (within-participants, unimodal/congruent/incongruent), TARGET (target/non-target; only used for response times, see below) and tACS (within-participants, sham/beta/gamma). First, we report detection accuracies using signal detection theory (SDT) measures d’ and c ^30, 31^. Second, we present the results of a distribution-level analysis of response times (RTs) based on cumulative distribution functions (CDFs)^32^. Concluding, we show correlations of sensory capability and behavioural performance with measures of cognitive status.

### Detection accuracies

To describe the accuracy of responses in our visuotactile task, we used SDT measures sensitivity d’ and criterion location c as a metric of response bias (see *Methods* for details). Sensitivity measure d’ and bias c were subjected to a mixed repeated-measures ANOVA design containing the factors AGE, STIMULUS, and tACS. In contrast to the RT analysis, the factor TARGET is missing from the analysis due to the way sensitivity measure d’ and bias c are calculated (see *Methods* and Figure 2). In the following, all ANOVA *p*-values are reported using Greenhouse-Geisser correction. Post-hoc analysis was done using (unpaired or paired) t-tests applying Bonferroni correction for multiple comparisons if appropriate. If not stated otherwise, all effects are reported significant at *p* < 0.05. In addition, we computed effect sizes as indicated by partial eta squared for ANOVAs 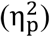, and Cohen’s *d* for pairwise comparisons.

**Figure 2.**
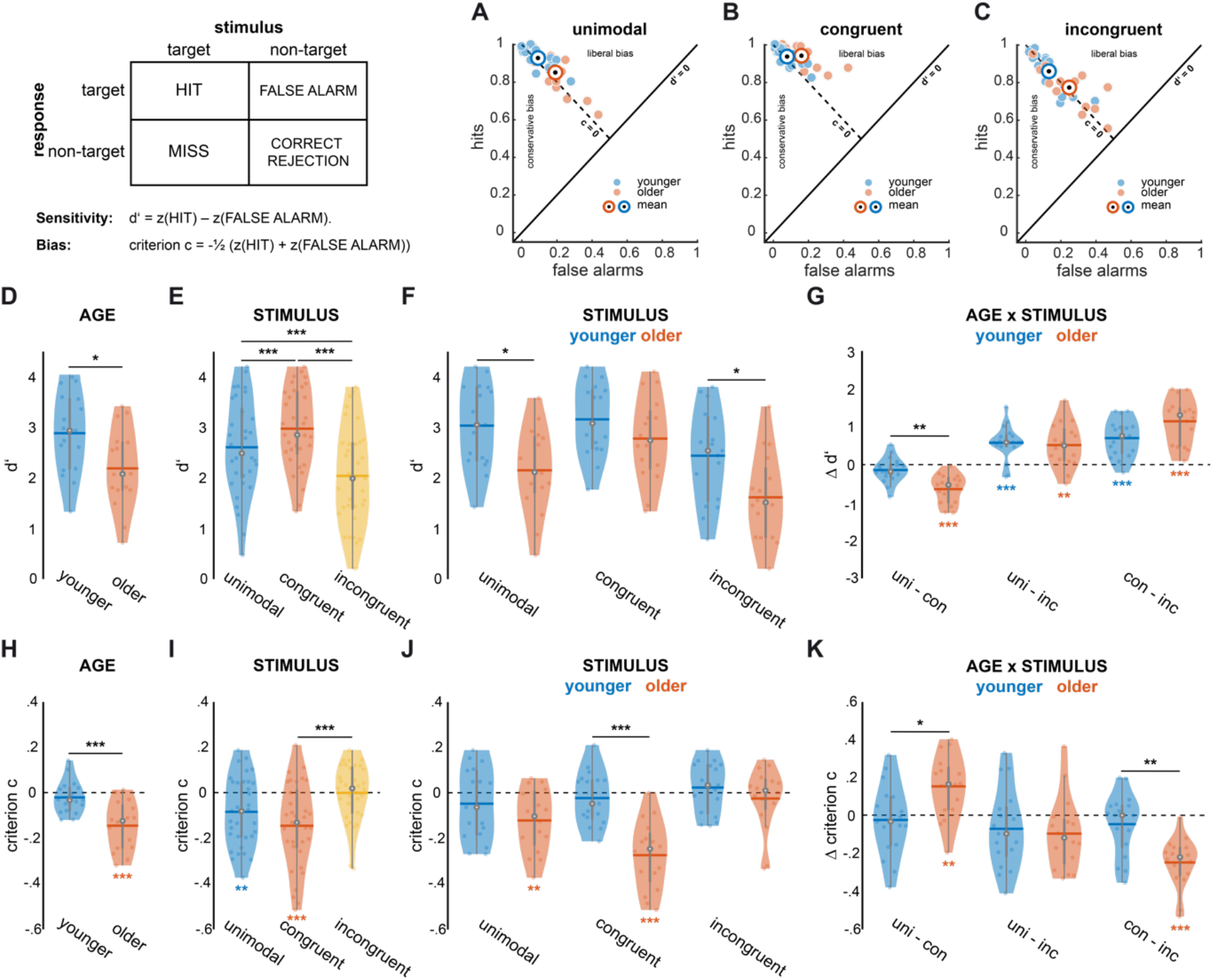
Age- and stimulus-related differences in sensitivity d’ and bias c. Violin plots show mean value (horizontal line), median (white dot), interquartile range (horizontal line) and data of individual participants (coloured dots). Black asterisks indicate statistical significance of t-tests for pairwise comparisons, coloured asterisks indicate significance for one-sample t-tests against zero (* *p* < 0.05, ** *p* < 0.01, *** *p* < 0.001). **(A)-(C)** Hits and false alarms as a function of STIMULUS, separately for younger and older groups. **(D)** Effect of AGE on d’. **(E)** Effect of STIMULUS on d’. **(F)** and **(G)** Interaction between AGE and STIMULUS for d’ scores is shown as difference between conditions of STIMULUS, separately for younger and older groups. **(H)** Effect of AGE on criterion c. **(I)** Effect of STIMULUS on criterion c. **(J)** and **(K)** Interaction between AGE and STIMULUS for criterion c. All conditions of STIMULUS are shown separately for younger and older groups.

For sensitivity measure d’, a main effect of AGE (*F*_1,31_ = 6.44, *p* = 0.016, 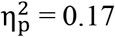 see Figure 2 D) was observed with younger adults showing significantly better overall performance (2.89 ± 0.79 vs. 2.19 ± 0.79, mean ± standard deviation (SD)). In addition, we found a main effect of STIMULUS (*F*_2,62_ = 62.61, *p* < 0.001, 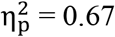 Figure 2 E) as well as an interaction of AGE and STIMULUS (*F*_2,62_ = 5.28, *p* = 0.012, 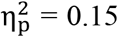; see also Figure 2 F, G). Tactile patterns appearing with congruent visual patterns were associated with the best performance (2.98 ± 0.79), followed by unimodal patterns (2.6 ± 0.94) and incongruent stimulus pairs (2.04 ± 0.97). Post-hoc analysis including Bonferroni correction (dividing the significance level α by 3 resulted in a corrected α of 0.0167) showed that all stimulus levels (unimodal, congruent and incongruent) differed significantly from each other (paired-samples t-tests: unimodal vs. congruent: *t*_32_ = −4.84, *d* = −0.84; unimodal vs. incongruent: *t*_32_ = 6.28, *d* = 1.09; congruent vs. incongruent: *t*_32_ = 9.14, *d* = 1.59; all *p*-values < 0.001). Post-hoc analysis of the interaction effect (including Bonferroni correction) showed that only the contrast of unimodal and congruent stimuli yielded significance between older and younger participants (*t*_31_ = −4.02, *p* < 0.001, *d* = −1.4; Figure 2 G). In other words, older adults profited significantly more from crossmodal stimulus congruence in their detection performance compared to the younger group. Finally, tACS did not influence sensitivity measure d’.

The analysis of bias c showed a similar overall pattern: Main effects of AGE (*F*_1.31_ = 15.4, *p* < 0.001, 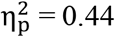; see Figure 2 H) and STIMULUS (*F*_2,62_ = 11.4, *p* < 0.001, 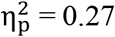; Figure 2 I) as well as a significant interaction of these two factors were observed (*F*_2,62_ = 6.32, *p* = 0.005, 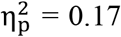; Figure 4 J, K). The average bias for the older group was −0.14 (SD = 0.11), indicating a liberal response tendency, whereas the younger group obtained an average value of −0.02 (SD = 0.07; see also Figure 2 H). Interestingly, only the older participant group showed a response bias significantly different from zero (*t*_15_ = −5.28, *p* < 0.001, *d* = −1.32). Average biases across all participants were the following: −0.08 ± 0.15 for the unimodal condition, −0.15 ± 0.18 for congruent stimulus pairs and −0.0008 ± 0.13 for incongruent pairs. Post hoc tests revealed that the stimulus effect was driven by the difference between congruent and incongruent conditions (*t*_32_ = −4.62, *p* < 0.001, *d* = −0.8; Figure 2 I) while all other contrasts were not significant. Of note, only the unimodal (*t*_32_ = −3.19, *p* = 0.003, *d* = −0.56) and the congruent (*t*_32_ = −4.54, *p* < 0.001, *d* = −0.79) conditions were associated with bias values significantly different from zero. Bias differed between older and younger participants between unimodal and congruent conditions (*t*_31_ = 2.86, *p* = 0.008, *d* = 1), as well as between congruent and incongruent stimulus pairs (*t*_31_ = −3.87, *p* < 0.001, *d* = −1.35; Figure 2 K). For both contrasts, differences between conditions were more pronounced for the older participant group. Finally, tACS did not influence bias c.

### Response times

CDFs were estimated as Gumbel functions using kernel density estimation^33^ on condition-pooled data. Pairwise comparisons reflecting main and interaction effects of all within- and between-participants factors of the mixed ANOVA design were performed by subtraction of CDFs. Non-parametric statistical evaluation was based on confidence intervals estimated by subtraction of shuffled data CDFs. Reported *p*-values reflect the minimum level of significance that needs to be exceeded along the whole range of RTs. Instead of exact *p*-values for the individual effects, we report RT ranges in which the *p*-value is undershot. Effect sizes were estimated for pairwise comparisons with Cohen’s *d* at the response time of the maximal group effect. Details of the statistical approach can be found in the *Methods* section.

First, we found main effects for the factors AGE, TARGET and STIMULUS (Figure 3). Overall, younger participants responded faster than older participants (Figure 3 A; CDF threshold: younger 787 ms, older 926 ms). This difference was significant across the whole range of RTs and showed a large effect size (*p* < 0.0013; younger-older: 0-2500 ms, *d* = 1.15 at 824 ms). Furthermore, target and non-target trials differed significantly in response speed. When the pattern to be detected was presented in the tactile channel (target), participants responded significantly faster compared to when a non-target would be presented as the tactile stimulus (Figure 3 B). This difference was significant for responses between 720 and 1720 ms and showed a large effect size at maximum (*p* < 0.00131, *d* = 0.78 at 1016 ms). Lastly, we found RT distributions to be significantly shaped by the nature of the stimulus. Unimodal stimuli showed overall fastest responses, congruent visuotactile stimuli were relatively delayed and incongruent visuotactile stimuli were slowest (Figure 3 C). All pairwise comparisons showed significant differences with medium to large effect sizes (*p* < 0.00039; unimodal-congruent: 830-1870 ms, *d* = 0.40 at 975 ms; unimodal-incongruent: 700-2120 ms, *d* = 0.95 at 993 ms; congruent-incongruent: 780-1540 ms, *d* = 0.77 at 1002 ms).

**Figure 3.**
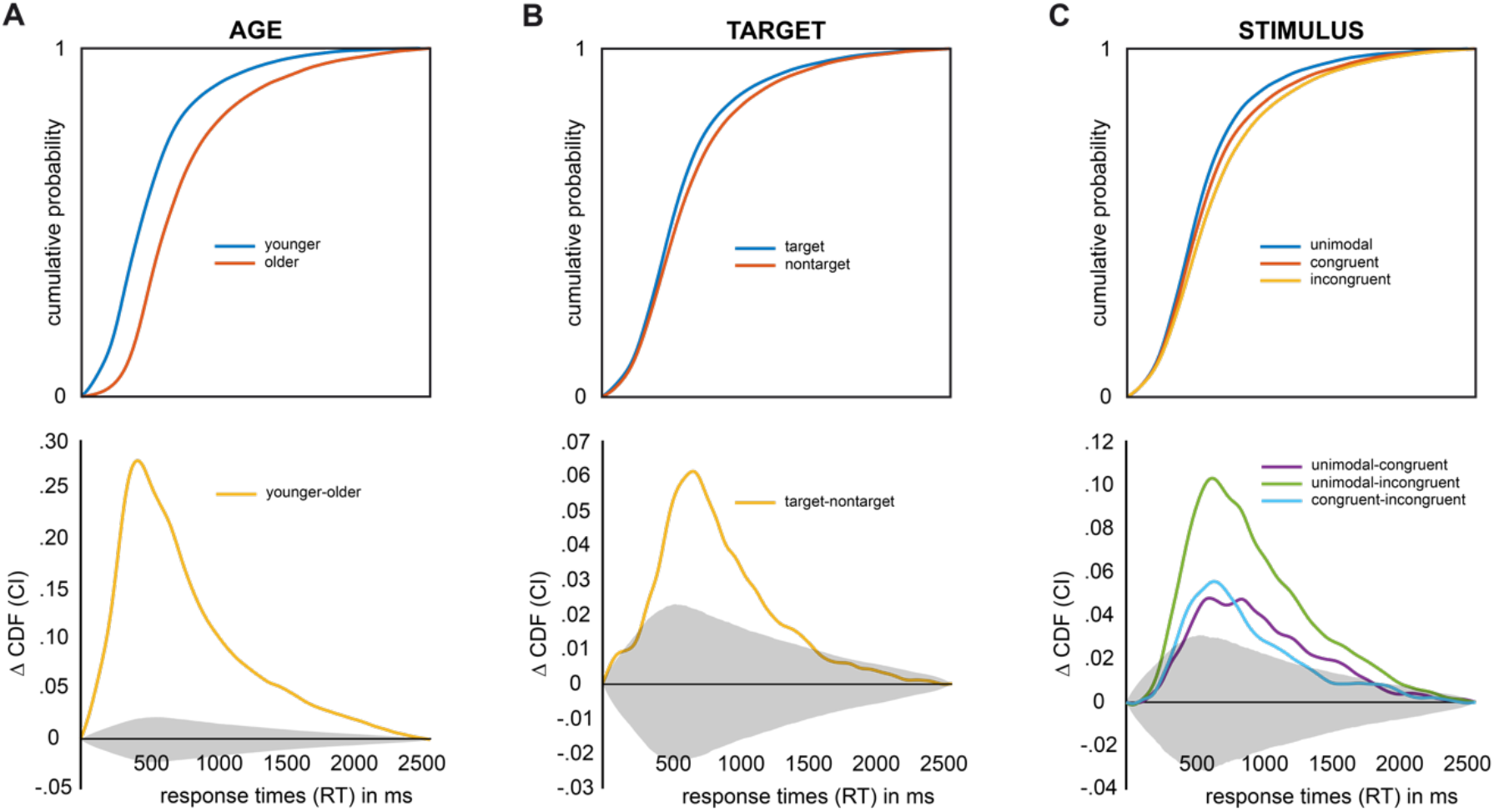
Main effects of AGE, TARGET and STIMULUS on cumulative distribution functions (CDFs) of response times (RTs). Grey shaded areas depict confidence intervals corrected for multiple testing. **(A) (top)** CDFs broken down by AGE. **(bottom)** Pairwise differences between AGE groups. **(B) (top)** CDFs broken down by TARGET. **(bottom)** Pairwise differences between TARGET conditions. **(C) (top)** CDFs broken down by STIMULUS. **(bottom)** Pairwise differences between STIMULUS conditions.

Secondly, the experimental factors interacted in shaping response times (Figure 4). AGE affected the amount of both the TARGET and the STIMULUS effect. Although both age groups showed significantly faster responses on target trials (Figure 4 A top; *p* < 0.00065; younger: 0-1610 ms, older: 720-1820 ms and 1860-2090 ms), older participants showed a significantly larger positive effect of target presentation (Figure 4 A bottom; *p* < 0.00145; younger-older: 910-1140 ms, *d* = 0.64 at 977 ms). Significant stimulus effects were seen in both age groups for all but one comparison: in younger participants, unimodal and congruent trials did not differ with respect to response times (Figure 4 B top; *p* < 0.00018, younger unimodal-congruent: none, younger unimodal-incongruent: 700-1300 ms, younger congruent-incongruent: 770-1160 ms, older unimodal-congruent: 850-2260 ms, older unimodal-incongruent: 700-2370 ms, older congruent-incongruent: 740-2090 ms). When comparing these simple effects of stimulus condition across age groups, we found that the difference between unimodal and congruent as well as incongruent conditions was significantly stronger in older participants (Figure 4 B bottom; *p* < 0.00039, younger-older, unimodal-congruent: 1010-2160 ms; younger-older, unimodal-incongruent: 990-2270 ms). This interaction showed large or very large effect sizes (unimodal-congruent: *d* = 1.21 at 1163 ms; unimodal-incongruent: 990-2270 ms, *d* = 1.05 at 1165 ms). The contrast between congruent and incongruent conditions, however, did not show significant difference with respect to age (Figure 4 B bottom). Additionally, stimulus effects differed between target and non-target trials. That is, the beneficial effect of the target was not present for incongruent stimulus presentations (Figure 4 C top; *p* < 0.00039, unimodal: 670-1480 ms, congruent: 500-630 and 760-2130 ms, incongruent: none). Pairwise comparisons of the target effect showed that the effect’s magnitude was comparable between unimodal and congruent stimuli, but was significantly smaller in incongruent trials compared to both others (Figure 4 C bottom; *p* < 0.00039; unimodal-congruent: none; unimodal-incongruent: 830-1090 ms, *d* = 1.00 at 938 ms; congruent-incongruent: 910-1360 ms, *d* = 0.72 at 1004 ms).

**Figure 4.**
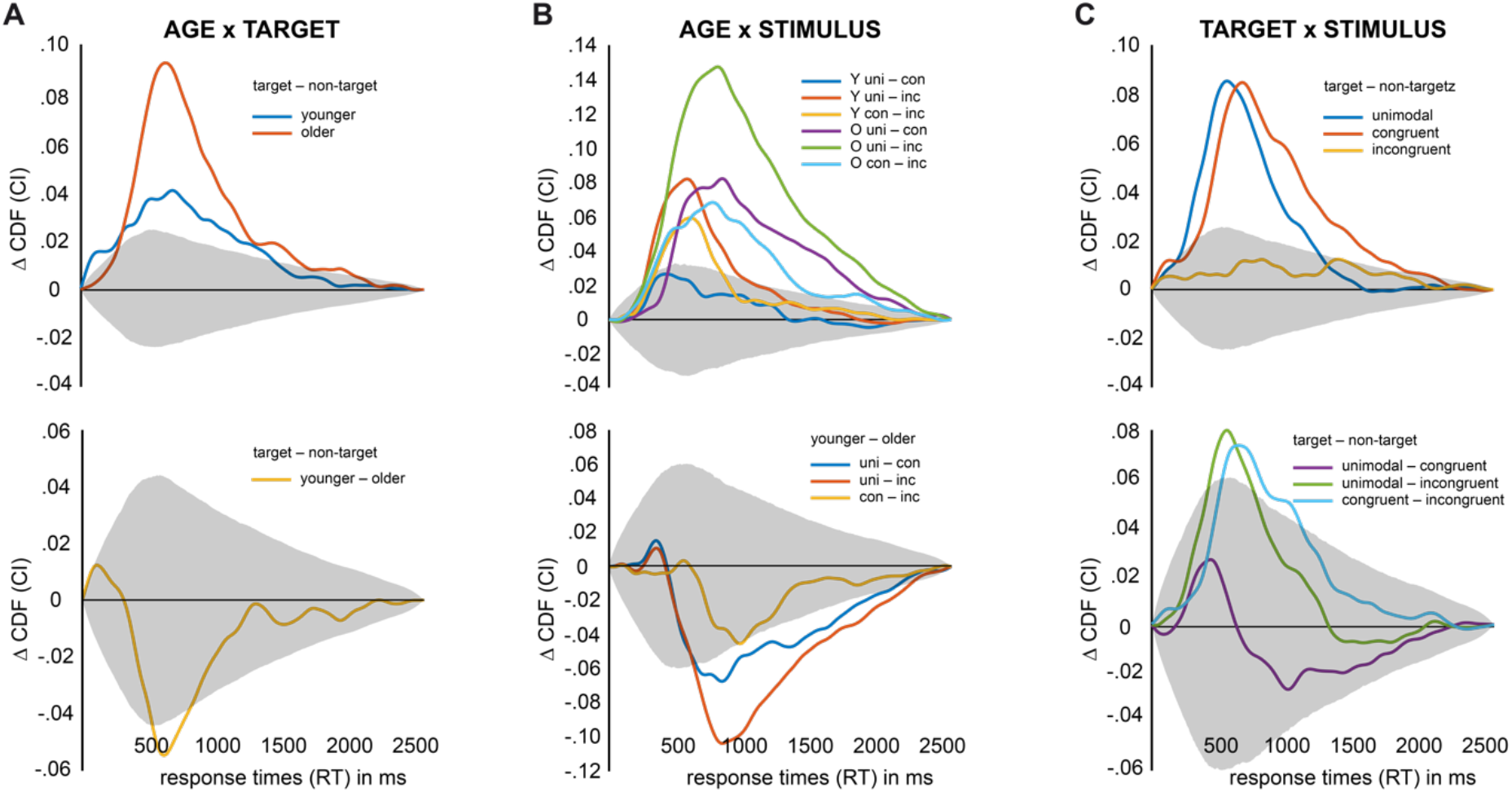
Interactions between experimental factors AGE, TARGET and STIMULUS on cumulative distribution functions (CDFs) of response times (RTs). Grey shaded areas depict confidence intervals corrected for multiple testing. **(A)** Interaction between AGE and TARGET. **(top)** Simple effects of TARGET separately for both AGE groups. **(bottom)** Differences between simple effects of TARGET. **(B)** Interaction between AGE and STIMULUS. **(top)** Simple effects of STIMULUS separately for both AGE groups. **(bottom)** Differences between simple effects of STIMULUS. **(C)** Interaction between TARGET and STIMULUS. **(top)** Simple effects of STIMULUS for both levels of TARGET. **(bottom)** Differences between simple effects of STIMULUS.

Finally, tACS influenced response times. Overall, we found that beta stimulation led to more fast responses and gamma stimulation to fewer fast responses, a result pattern that was highly comparable between age groups (Figure 5 A-C). Yet, due to the strong main effect of AGE, tACS effects of the older group were shifted rightwards on the x axis. As a consequence, comparing tACS effects between age groups resulted in a marginally significant interaction (see Supplementary Figure S1 C). In order to account for the main effect of age, we normalized CDFs with the threshold of the age groups’ average CDF (Supplementary Figure S1 D, E; threshold younger: 787 ms, threshold older: 926 ms), resulting in a non-significant interaction between AGE group and tACS condition (Supplementary Figure S1 F). The tACS effect based on normalized CDFs is depicted in Figure 5 D (*p* < 0.00028). Under beta stimulation, the likelihood of fast responses between −230 and −110 ms relative to the group median was significantly increased compared with sham. Gamma stimulation, in contrast, led to a significant decrease of fast responses when compared with sham (−110 to 10 ms relative to the group median). As a consequence, we found significant differences between beta and gamma stimulation for responses between −230 and −20 ms relative to the group median. Effect sizes were small or very small, respectively (sham-beta: *d* = 0.11 at −142 ms; sham-gamma: *d* = 0.16 at −46 ms; beta-gamma: *d* = 0.15 at −82 ms). Importantly, we found that tACS effects were only significant for target trials (Figure 5 E-F). While the slowing of responses under gamma tACS was equally pronounced for younger and older participants, the speeding effect of beta tACS was significantly more pronounced in the older group (Figure 5 G-I).

**Figure 5.**
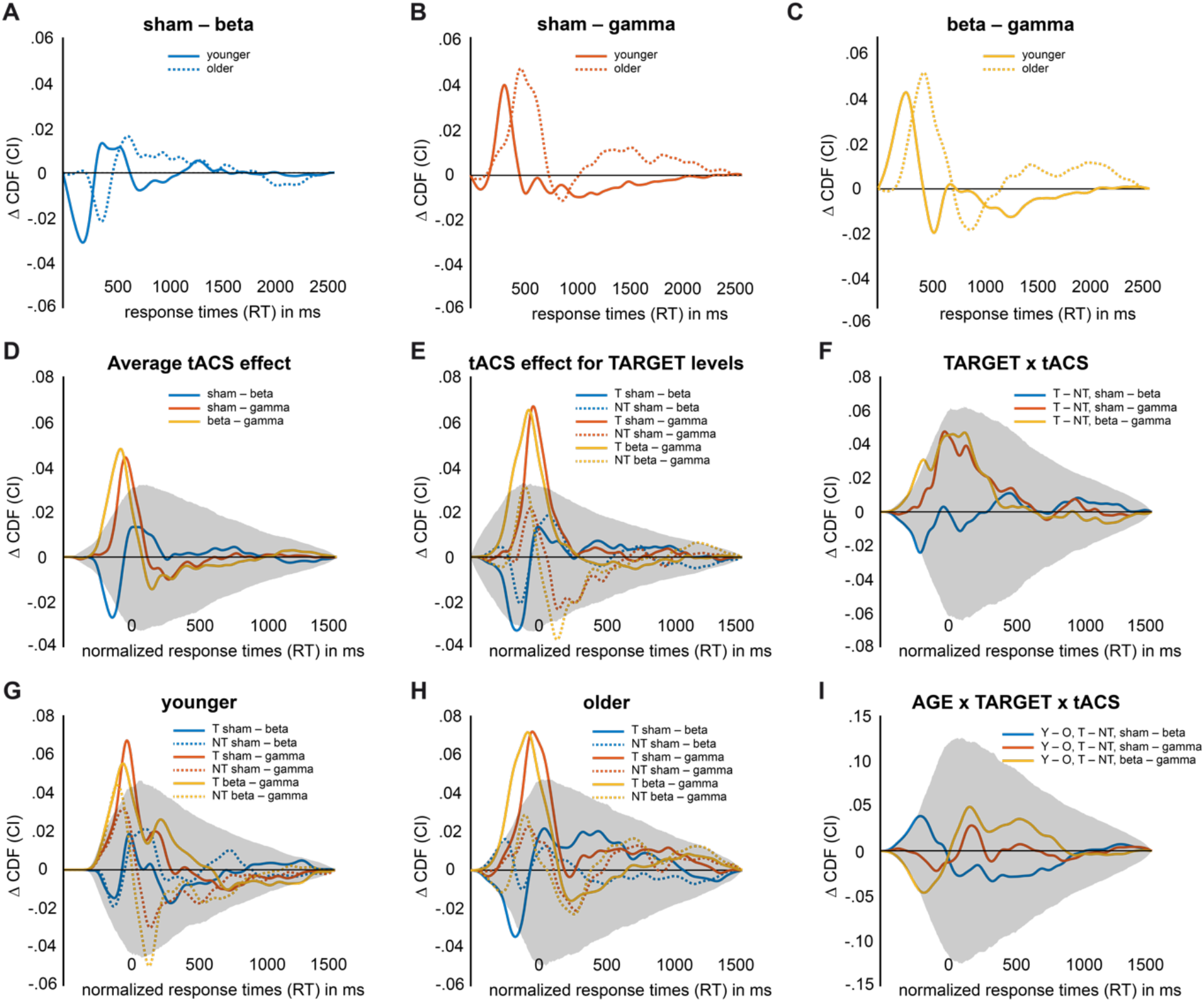
tACS-related differences between cumulative distribution functions (CDFs) of response times (RTs). Grey shaded areas depict confidence intervals corrected for multiple testing. **(A)** Effect of Beta tACS (20 Hz) on uncorrected CDFs shown separately for age groups. **(B)** Effect of Gamma tACS (70 Hz) on uncorrected CDFs shown separately for age groups. **(C)** Difference between Beta and Gamma tACS on uncorrected CDFs shown separately for age groups. **(D)** Average tACS effect on normalized CDFs. **(E)** tACS effect on normalized CDFs shown for target (solid lines) and non-target (dashed lines) trials. **(F)** Interaction effect between TARGET and tACS. **(G)** The same as in B, but shown only for the younger group. **(H)** The same as in B, but shown only for the older group. **(I)** Interaction between AGE, TARGET and tACS.

### Relation between behavioural performance and cognitive status

In order to integrate our experimental findings with the assessment data, we applied linear correlation analysis to quantify how measures of sensory capability and behavioural performance relate to the cognitive status of our participants. Bivariate correlations were computed between the following variables: DemTect scores, age, grand average RTs, and response bias (grand average of criterion c). Additionally, we included sensitivity d’ and sensory acuity (a composite measure for visual and tactile thresholds; see *Methods* for details) as estimates of sensory capability. Finally, two more metrics entered the correlation analysis: the difference between d’ values for congruent and unimodal conditions, termed multisensory congruence enhancement (MCE)^14, 34^ as well as the target effect found in our RT data, labelled working memory enhancement (WME; CDF RT difference between targets and non-targets; see *Methods* for details). In the following, we consider MCE and WME as indices of information integration, either mediated by crossmodal stimulus congruence or working memory benefits.

We refrained from using MMSE scores as an index of cognitive status because neither younger nor older participants showed sufficient variability in this measure. DemTect scores in contrast were more sensitive for inter-individual differences in the older group, but again did not show sufficient variability in the younger group. We thus restricted this analysis to the older group. Age within the group of older participants did not correlate significantly with any of the other variables, but showed a trend towards longer RTs for higher age (*r* = 0.471, *p* = 0.065). Similarly, overall bias did not predict cognitive status or any of the other variables. Of the remaining variables, we found two groups that independently predicted cognitive status. On the one hand, estimates of sensory capability (sensitivity d’ and sensory acuity) were positively correlated (*r* = 0.650, *p* = 0.006; see Figure 6 A) and predicted DemTect such that higher cognitive status was related to higher levels of sensory capability (*r* = 0.584/0.604, *p* = 0.0180/0.0133; see Figure 6 A). On the other hand, WME and MCE (as estimates of information integration) were positively correlated (*r* = 0.655, *p* = 0.006; see Figure 6 A) and predicted DemTect such that lower cognitive status was related to enhanced integration (*r* = −0.686/−0.612, *p* = 0.003/0.012; see Figure 6 A). Importantly, we did not observe significant correlations between estimates of sensory capability and estimates of information integration (all |*r*| < 0.4 and all *p* > 0.14). Using partial correlations, we confirmed that both groups of predictors explained largely independent parts of the variance in DemTect scores (see Figure 6 B, C). Each of the correlations stated above remained significant even when controlling for both other variables of the other group. Of note, controlling for the influence of MCE and WME, we found an additional correlation between RT and age such that higher age was associated with slower responses (*r* = 0.636, *p* = 0.0144).

**Figure 6.**
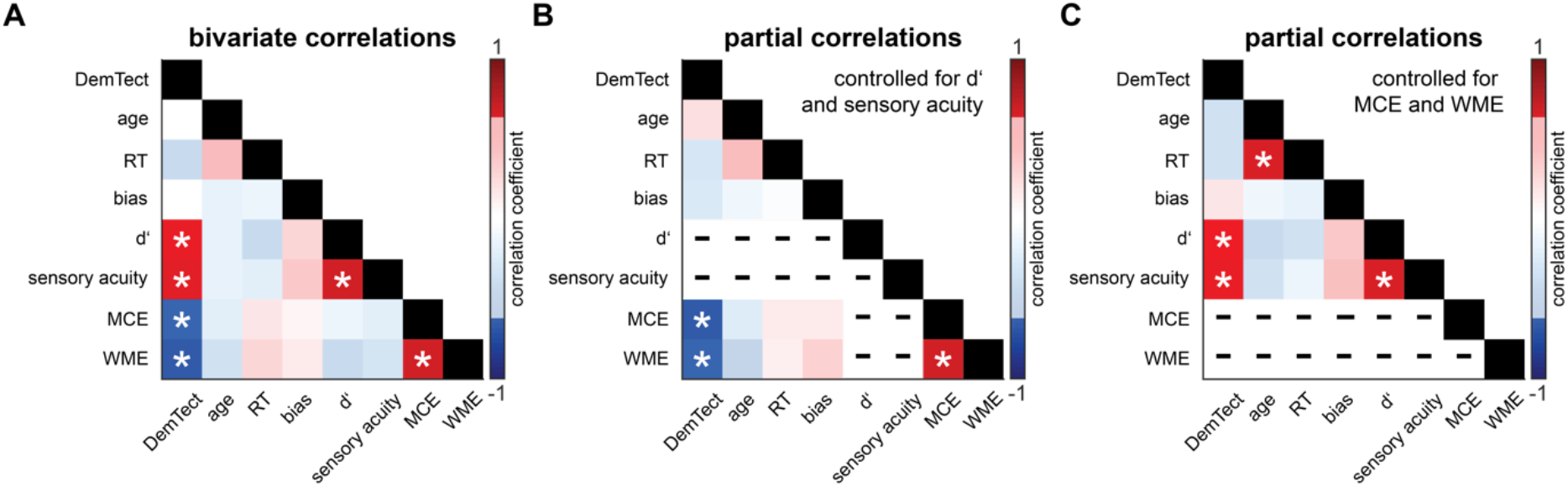
Correlations of cognitive status with perceptual and behavioural variables. Correlation coefficients are thresholded at *p* < 0.05 and non-significant correlations are shaded. Auto-correlations are marked with black and correlations that were partialled out are marked with minus signs. Significant correlations are marked with a white asterisk. **(A)** All pairwise bivariate correlations. **(B)** Partial correlations controlled for d’ and sensory acuity. **(C)** Partial correlations controlled for MCE (multisensory congruence enhancement) and WME (working memory enhancement).

### tACS-related adverse events

None of the participants reported severe or moderate adverse effects^35^ related to the electrical stimulation. Mild adverse effects^35^ were monitored using a custom-made questionnaire that quantified the intensity and time course of skin sensations, phosphenes and pain (see *Methods*). Older participants did not report significant mild adverse events related to stimulation (Figure S2 C; median intensities: skin sensations 0±0, phosphenes 0±0, fatigue 0±0, pain 0±0; all *p* > .5). Younger participants reported skin sensations (1±0.75, *p* = 0.001) and fatigue (1±2, *p* = 0.004) but no phosphenes (0±0, *p* = 0.5) or pain (0±0, *p* = 0.25). None of the comparisons within groups were significant between conditions (all *p* > 0.1). Between-group comparisons showed that younger participants perceived significantly stronger skin sensations (*p* = 0.002) and fatigue (*p* = 0.026) compared with the older group (Figure S2 A). We additionally asked to rate the time course of sensation. Ratings across all sensations were used to compute binary variables indicating sensations only in the beginning (0) or at any or all later time points (1; mean values younger: sham .24, beta .82, gamma .47; mean values older: sham .06, beta .31, gamma .13). Chi-square tests showed that younger participants were not significantly blinded for the difference between sham and beta-/gamma-frequency stimulation (*p* = 0.001), but comparisons between beta- and gamma-frequency were non-significant (Figure S2 B). Older adults were blinded for all comparisons between sham, gamma- and beta-frequency stimulation (Figure S2 D; all *p* > .2).

## Discussion

In our match-to-sample task, irrelevant visual input delayed responses overall, but showed differential effects on tactile detection sensitivity: while crossmodal incongruence of tactile and visual patterns was associated with diminished discrimination sensitivity, crossmodal congruence improved detection. Both costs and benefits of visuotactile interactions were more pronounced in the older group. At the same time, however, older participants showed a liberal response bias, which was absent in the younger group. Further, all participants showed speeded responses when the tactile target pattern was presented. Again, this effect was stronger in the older group. tACS with beta frequency (20 Hz) speeded responses whereas gamma stimulation (70 Hz) delayed responses. Strikingly, the speeding effect of beta-frequency stimulation was more pronounced in the older group, but only for target stimuli. Finally, we found that measures for sensory capability were independent of behavioural benefits of multisensory integration and working memory match, while both predicted cognitive status.

Numerous studies have demonstrated that multisensory perception changes across the lifespan^36, 37^. Our data adds to these findings and supports the hypothesis that ageing is characterized by enhanced multisensory interactions in tasks with redundant crossmodal input^12^. Specifically, older adults showed increased sensitivity in discriminating tactile dot patterns due to task-irrelevant but congruent visual co-stimulation, an effect that was absent in younger participants. This multisensory enhancement of tactile stimulus processing led to a levelling of discrimination sensitivity between younger and older adults. Potential for compensation of decreased sensory acuity by enhanced multisensory integration has been shown before^9, 17^ and was suspected to follow directly from the degradation of sensory systems^36^. If this was the case, we should thus observe a correlation between measures of sensory acuity and the extent of multisensory enhancement, which was not the case (Figure 7). Therefore, our data suggests that, while older participants can compensate loss in sensory acuity through multisensory integration to some degree, enhanced integration in ageing does not simply follow from the degradation of sensory systems.

**Figure 7.**
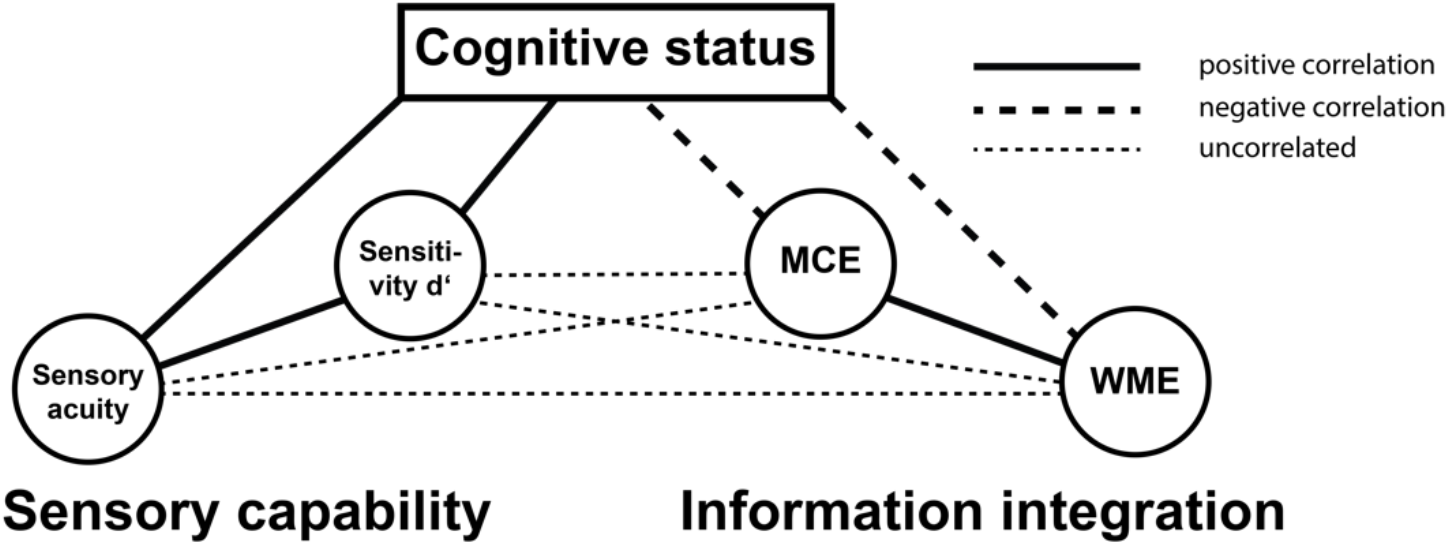
Two independent predictors of cognitive status. Lines depict linear correlations between variables (solid: positively correlated; dashed: negatively correlated; dotted: uncorrelated). Sensory acuity is a composite measure of thresholds for tactile and visual stimuli. Sensitivity d’ is the grand average from the signal detection analysis. MCE (multisensory congruence enhancement) is the simple effect of increase in sensitivity d’ from unimodal to congruent trials (congruent – unimodal). WME (working memory enhancement) is the response time effect of target versus non-target trials (non-targets – targets).

Importantly, however, we found the beneficial multisensory effect on detection sensitivity in the older group to be confounded by a liberal response bias. Older adults showed a tendency to report detection of the target pattern even in its absence and thus produced more false alarms. Specifically, we found a liberal decision criterion in unimodal trials, which was enhanced for congruent trials but absent for incongruent ones. Similar results were reported by Mishra and Gazzaley^38^ who described crossmodally enhanced sensitivity in older adults to be confounded by elevated false alarm rates in an audiovisual task. The authors contributed this finding to age-related deficits in inhibitory control^3, 39^, which in turn have been associated with decreased structural integrity of the prefrontal cortex^39, 40^. Although there is a lack of studies investigating response tendencies as a function of age in crossmodal settings, a rich body of literature describes bias differences related to age in the context of recognition memory^41–43^. It has been proposed that these alterations in decisional processes result from false attribution of familiarity^44, 45^ and behavioural studies support the idea of an age-related shift from recollection-based to familiarity-based mnemonic strategies^41,43,46^. Interestingly, familiarity has also been shown to play an important role in audiovisual integration^47, 48^. It thus should be considered that modulations of response bias due to multisensory characteristics of the stimuli might be an overlooked finding in other studies that evaluated overall response accuracy. As a consequence, the magnitude of multisensory enhancements might have been overestimated.

In the current study, response times to multisensory stimuli were overall delayed compared with unimodal stimuli. Slowing of responses was especially pronounced for crossmodally incongruent information^49^, but also present for congruent visual patterns. This finding contrasts with studies showing speeded responses to redundant/congruent crossmodal stimuli in simple reaction time tasks^50, 51^ or when evaluation of crossmodal congruence was explicitly instructed^14,52–55^, but is well compatible with studies that report crossmodal distraction in working memory paradigms^56, 57^. In our task, tactile patterns had to be matched with a sample pattern held in working memory and, thus, our data supports the notion that working memory processes are vulnerable to crossmodal distraction. Importantly, we showed that this effect was enhanced in the older group, which is line with earlier findings of increased distractibility of older adults in a visuotactile task^58^. On the other hand, we report marked improvements of response speed for tactile target patterns, that is, when matching between working memory content and sensory information was successful. This finding is in line with studies showing enhanced processing of sensory stimuli that match the content of working memory^59–61^. Notably, this effect was moderate for younger adults but strongly enhanced in the older group. Taken together, we present evidence for older adults’ increased crossmodal distractibility^40, 62^ as well as their enhanced benefit of successful working memory matching.

In a correlation analysis (see Figure 6 and Figure 7), we found that the target effect in response speed (WME) was tightly linked to the congruence-mediated effect in sensitivity (MCE). Substantial evidence suggests that both multisensory integration and working memory match rely on information integration that is probably implemented by interactions of oscillatory activity in cortical networks^25,54,55,63,64^. We speculate that both the target effect in RT as well as the congruence-related benefit in sensitivity found in the older group reflect the underlying tendency of enhanced information integration, possibly related to altered oscillatory coupling between involved cortical regions. A candidate structure that has been highlighted as a hub in multisensory as well as working memory networks is the parietal cortex^65–67^. A central role of the parietal cortex in mediating enhanced information integration in our older adult group is partly supported by the effects of tACS. Compared with sham, 20 Hz tACS over left IPS and S1 speeded responses while 70 Hz stimulation delayed responses in our study. Importantly, the effects of beta-frequency tACS were significant only for target trials in the older group, corroborating that older adults rely more on the integrative functions of the parietal cortex during working memory matching. Specifically, we propose that boosting beta power in a somatosensory-parietal network facilitated the reactivation of working memory content and possibly also supported matching the reactivated pattern and the sensory information in somatosensory cortex^68^. Gamma-frequency stimulation, on the other hand, might have had an overall inhibiting impact on beta oscillations and, thus, hindered reactivation and/or working memory matching. This opposing nature of beta- and gamma-frequency tACS has been noted earlier^69–72^, and has been suggested to reflect cross-frequency modulations between gamma- and beta-band oscillations^72, 73^. Taken together, we propose that older adults generally exhibit a higher tendency of information integration that might be related to an altered role of parietal oscillatory activity in cortical network dynamics.

Assessing the relation of participants’ behaviour in our crossmodal task and DemTect scores (see Figure 6 and Figure 7), we replicated earlier findings that showed the predictive value of sensory capability^74^ and multisensory integration^13^ for cognitive status in healthy older adults. Importantly, we add to the existing literature by showing that these two factors are independent and likely represent two separate processes of cognitive ageing that in turn might result from distinct processes of brain ageing. On the one hand, a decline in sensory capability might result from degraded integrity of sensory organs and/or peripheral nerves. Of note, artificial degradation of sensory information in younger adults has been shown to be insufficient to produce a reduction in cognitive performance, indicating that the correlation between sensory and cognitive factors in older adults might merely be epiphenomenal^75^. On the other hand, we show that enhanced multisensory integration correlates positively with working-memory matching and, thus, might reflect an overall enhanced tendency of older adults to integrate information linked to altered function of the parietal cortex. Although common theories of cognitive ageing do not specifically capitalize on the function of the parietal cortex, many ideas postulate a special role of the prefrontal cortex^5,39,76^. For most cognitive functions, however, (pre-) frontal cortex does not achieve cognitive control in unitary action but operates tightly coupled to the parietal cortex^67, 77^. We propose that in the current study decreased integrity of the frontal cortex might not only be the substrate for the enhanced bias observed in older adults^38, 41^ but, as a secondary effect, lead to altered parietal dynamics. Specifically, we speculate that enhanced (bottom-up) information integration in ageing might result from a deficiency of frontal cortex to exert top-down modulation over posterior cortex. Bearing this speculation in mind, future studies are advised to broaden the focus from structural and functional changes in frontal cortex to potential alterations in parietal brain areas.

Implications of our results are limited in a number of important ways. First, the lack of a structural magnetic resonance imaging (MRI) analysis prevents definite conclusions about the impact of prefrontal integrity on cognitive status or its predictors, respectively. Furthermore, not using individual MR images as head models prevented us from adapting tACS montages for optimal targeting of cortical sources. Also, we designed montages based on a task-positive beta network found in an earlier study in younger adults^25^ and, thus, did not account for potential age-related changes in location and frequency of the underlying networks. Yet, finding significant modulation by tACS that was largely comparable across age groups suggests that this approach can be successful. Nonetheless, individual tailoring of stimulation parameters will most likely reduce variance and elevate effect sizes. Finally, we did not record M/EEG and thus cannot detail on the functional underpinnings of the behavioural effects.

In conclusion, we show that age-related changes in multisensory integration do not result from sensory degradation. We argue that enhanced multisensory integration in older adults likely reflects an overall enhanced tendency to integrate information that, in our data, was also reflected by behavioural benefits of successful working memory matching. A central role of the parietal cortex in mediating augmented integration in age is supported by effects of beta-frequency tACS found in the current study. Importantly, indices of sensory capability and information integration predicted cognitive status in the absence of disease. We suggest that these two independent groups of predictors of cognitive status result from distinct processes of brain ageing that should be studied separately.

## Methods

### Participants

We re-invited 13 younger and 15 older adults from a cohort that was enrolled in a previous behavioural study^15^ and additionally recruited new participants in order to include 17 individuals in each group. In the older group, we excluded one re-invited participant due to a decline in the cognitive status (DemTect < 13). The final sample consisted of 17 younger (10 female, 24.77±3.09 years) and 16 older (10 female, 72.56±4.55 years) volunteers. All participants were right-handed according to the Edinburgh handedness inventory^78^, had age-adjusted normal or corrected-to-normal vision, no history or symptoms of neuro-psychiatric disorders and no history of centrally acting drug intake. All participants received monetary compensation for participation in the study. The study was conducted in accordance with the Declaration of Helsinki and was approved by the local ethics committee of the Medical Association of Hamburg (PV5085). All participants gave written informed consent.

### Assessment

Prior to the experiment, every participant underwent an assessment procedure consisting of a neurological examination, the Mini-Mental State Examination (MMSE)^28^ and the DemTect^29^ to rule out symptoms of neuro-psychiatric disorders and to examine the cognitive status. A 2-point-discrimination test^26^ was conducted to ensure intact peripheral somatosensation and a test of the visual acuity (Snellen chart)^27^ to ensure intact vision. Additionally, every participant rated the subjectively experienced attention level, general fatigue and fatigue of the hand at the beginning of the experiment by placing a mark on a 10-cm visual analogue scale (VAS). The extremes of the horizontally positioned VAS were labelled: attentive vs. inattentive, awake vs. tired, hand not tired vs. hand tired. The VAS values were analysed by measuring the position of the mark in cm.

### Experimental procedure

After the assessment, we prepared an EEG cap in which electrodes for the electric stimulation were mounted and applied an analgesic creme (see *tACS stimulation* for details). In a training session, participants got acquainted with the experimental setup, the stimulus material and experimental task. After the four dot patterns (Figure 1 B) were introduced visually and tactilely, participants were trained to match two consecutively presented tactile dot patterns. Training blocks of 24 pairs had to be completed until matching accuracy was at 75% correct or above (min. 3 blocks, max. 10 blocks). The experiment consisted of three main blocks of 17.5 minutes duration in which tACS was applied with sham, beta frequency (20 Hz) or gamma frequency (70 Hz) stimulation. The sequence of tACS conditions was counterbalanced across participants. At the beginning of each block, tACS was ramped up before experimental trials began (see *tACS stimulation* for details). Each block consisted of 192 trials, split in four sessions of 48 trials separated by short breaks of 30 s. After completion of each of the three main blocks, we asked participants to complete a questionnaire assessing the side-effects of the electric stimulation. Taken together, participants completed 576 trials in about 60 minutes.

### Experimental design

Participants’ task was to compare tactile patterns to a target pattern and report match or mismatch as fast and accurately as possible via button press (see also *Experimental setup*). The target pattern was chosen pseudo-randomly from the stimulus set (see Figure 1 B) and was newly defined for each of the four sessions within a block (so each of the four patterns served as target in every tACS block). Each trial started with a central fixation of 1000 ms, followed by the presentation of a unimodal tactile or crossmodal visuotactile stimulus for 500 ms (Figure 1 A). The tactile pattern was administered via a Braille stimulator (see *Stimulus material* for details), either unimodally (1/3 of all trials), or accompanied by a congruent (1/3) or incongruent (1/3) visual stimulus presented on a computer screen. The response interval was limited to 2750 ms, after which feedback was given by colouring the fixation cross green or red. In half of all trials in a given session of 48 trials, we presented the target pattern as the tactile stimulus. Overall, each block contained equally many stimuli of each condition (unimodal, congruent and incongruent) and featured all four patterns as targets in pseudo-randomized order.

### Experimental setup

The experiment took place in a light-attenuated chamber and participants were comfortably seated in a chair with their right hand resting on a custom-made board containing the Braille stimulator (QuaeroSys Medical Devices, Schotten, Germany). Tactile patterns were administered to the participants’ right index fingertip while visual patterns were presented on a 21-inch computer screen running at 60 Hz with a resolution of 1024 x 768 pixels, positioned 110 cm in front of the participants. Participants responded using their left middle or index finger (counterbalanced within both age groups) to press one of two buttons on a response box (Cedrus, Model RB-834, San Pedro, USA). We used Presentation software (Neurobehavioral Systems, Version 16.3) to control stimulus presentation and to record participants’ response times (RTs) and accuracies.

### Stimulus material

The Braille stimulation cell consisted of eight independently controllable pins arranged in a four-by-two matrix, each pin 1 mm in diameter with a spacing of 2.5 mm. The set of tactile stimuli used in the current experiment included four geometric patterns (see Figure 1B), each of them formed by four elevated pins. Visual stimuli were designed analogously to the tactile patterns and presented left of a central fixation cross on a noisy background (Perlin noise; Figure 1A). The visual patterns subtended 3.5° x 2.5° of visual angle. To achieve comparable performance in pattern detection in older and younger participants, tactile and visual stimulus intensities were adjusted individually. We matched the amplitude of pin elevation as well as the grey intensity of the visual patterns to individual thresholds^15^. The Braille stimulator used allowed the pin elevation to be controlled in 4095 discrete steps, with a maximum amplitude of 1.5 mm. Maximal grey intensity equalled black (RGB: 0-0-0). For participants who already participated in the preceding study, we used the originally estimated thresholds. The newly recruited participants underwent the thresholding procedure one week prior to the experimental session.

### tACS stimulation

Multi-electrode transcranial alternating current stimulation (tACS) was administered via two separate stimulators (DC-Stimulator Plus, Neuroconn, Germany) used in external mode. That means, current output was precisely controlled via voltage input to the stimulator that was produced by a NI-DAQ device run with Labview (NI USB 6343, National Instruments, USA). For each montage, we used five Ag/AgCl ring electrodes (diameter = 12 mm) organised in 4- in-1 montages. These montages aim to restrict current flow under the central electrode and thereby increase focality of stimulation^79^. To that end, electrodes were prepared such that impedances were homogenous within montages and as low as possible (below 50 kΩ). Cortical targets were chosen from a previous study that identified a left-hemispheric beta-band network to be associated with target detection in a similar paradigm as employed here^25^. In accordance with these results, we prepared montages over left primary somatosensory cortex (S1) and left intraparietal sulcus (IPS) and simulated resulting electric fields. Current densities were estimated by using the inverse model constructed by means of exact low-resolution electromagnetic tomography (eLORETA)80 on the basis of a boundary-element three-shell head model^81^ in combination with a cortical grid (MNI152). The resulting electric field at location 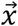 was then modelled as the sum of linear combinations between eLORETA leadfield 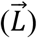 and injected currents at the stimulation electrodes (α_*i*_):

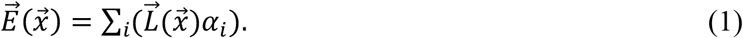

In order to compensate for differences in the anatomy of the skull (thickness, curvature, etc.) that impact the strength and depth of resulting electric fields, we determined the necessary amount of electric current to be administered over S1 and IPS in order to achieve current densities of up to 0.3 V/m in the targeted cortical regions. As a consequence, we applied 1.68 mA peak-to-peak over S1 and 2.4 mA peak-to-peak over IPS. Stimulation current was ramped up from 0 to maximal intensity within 10 s prior to the experimental blocks and afterwards delivered constantly with either 20 Hz (beta) or 70 Hz (gamma). In sham blocks, stimulation was terminated after the ramp-in. In order to minimize transcutaneous side-effects of stimulation, we used a creme of eutectic mixture of local anaesthetics (EMLA) applied to the skin under each ring electrode. Side-effects were tracked with a questionnaire after each tACS block.

### Statistical analyses

Statistical analyses were performed in Matlab (Version 9.1, MathWorks, Natick, MA, 2014) and RStudio (Version 3.5.4, R Core Team, 2017) using custom-made scripts.

### Assessment data

To test for group differences in the assessment data a multivariate analysis of variance (MANOVA, R’s *manova()* command) was performed with the values for 2-point-discrimination, visual acuity, tactile threshold, visual threshold, MMSE, DemTect, fatigue, attention level and fatigue of hand as dependent variables and group (younger participants vs. older participants) as the independent variable. For post-hoc analysis, two-sample t-tests were performed. As the results of the assessment parameters are not independent, Benjamini-Yekutieli correction^82^ was used to adjust for multiple comparisons.

### Response times

Response times (RTs) were analysed on the level of distributions. The shape of RT distributions between 500 and 2500 ms after stimulus presentation was estimated with Gaussian kernels^33^ and compared between conditions as cumulative distribution functions (CDFs). Statistical evaluation was performed with permutation statistics for the complete factorial design of *AGE* (younger/older) * *TARGET* (target/nontarget) * *STIMULUS* (unimodal/congruent/incongruent) * *TACS* (sham/beta/gamma). To that end, we shuffled single-trial data across conditions and participants in order to form N collections of RT data, where N corresponds to the maximal number of collections for computing a given contrast. For instance, estimating the null distribution for the evaluation of *TARGET*, we iteratively build two collections from the shuffled RT data, computed CDFs for each collection and subtracted the CDFs. The null distribution for each contrast was computed by storing the CDF differences of each permutation (100.000 permutations). Confidence intervals for each main factor and interaction were built by finding percentiles according to *alpha* and 100-*alpha*. Initially, *alpha* was set to 2.5/n %, where n is the number of minimal multiple comparisons of a given factor/interaction. This *alpha* accounts for both the two-tailed analysis and the multiple comparisons due to the factorial design, but not for the range of RTs tested. In other words, the type I error is controlled for locally at each RT bin, but not globally for the whole distribution. This was achieved by increasing percentiles until maximally *alpha* % of all null-differences would exceed the CI globally. The resulting CI controls type I error globally at *alpha*. The actual condition differences were then computed in a similar fashion by first pooling data across all participants and sub-conditions. CDFs were computed on the pooled data and subtracted according to the factorial design. Differences exceeding the CI at any point of the RT range were treated as a significant condition difference.

### Detection accuracy

Detection performance was analysed using measures provided by signal detection theory (SDT)^30, 31^, namely sensitivity d’ and criterion location c (or bias). Whereas sensitivity captures the perceiver’s ability to discriminate different choice alternatives (e.g., targets vs. non-targets or, more general, signal vs. noise), response bias describes the perceiver’s tendency to categorize stimuli as either the one or the other^83^, regardless of actual stimulus presence. In the case of target detection, bias statistics reflects the degree to which “yes” or “no” choices are favoured^31^. Both d’ and c are calculated from hit and false alarm rates. In our paradigm, we defined hits and false alarms for the different conditions as follows: hits were equated with correctly identified tactile targets, either presented without a synchronous visual pattern (*unimodal*), or in combination with a *congruent* respectively an *incongruent* one. False alarms in contrast are usually defined as 1 − correct rejections, in our case 1 − the ratio of correctly identified non-targets, again either appearing in their *unimodal*, *congruent* or *incongruent* version. For these three conditions, we computed d’ values by subtracting the z-transformed false alarm (FA) rates from the z-transformed hit (H) rates, within both older and younger adults and for the three tACS conditions separately:

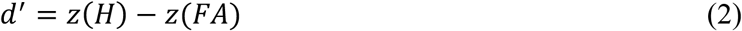

In the framework of SDT, bias measure c corresponds to the position of the decision criterion relative to the point of intersection between the (internal) distributions of signal and noise (or two signals). As in yes/no detection tasks, we computed c as:

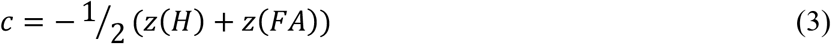

Response bias can be described as liberal (c < 0, corresponding to a tendency to say “yes” or in our case to (over-) detect targets), conservative (c > 0, corresponding to a tendency to detect non-targets) or neutral (c = 0, not systematically favouring either of the choice alternatives). In the terminology of SDT, a negative criterion in our study reflects a liberal bias to report targets for both trials where targets really are presented (hits) as well as non-target trials (false alarms). Analogously, a positive criterion value points to a conservative bias and a tendency to report non-targets in trials actually containing targets (misses) as well as in non-target trials (correct rejections). Lastly, a neutral bias is present if false alarm and miss rates are equal. To analyse task performance in our crossmodal detection paradigm, we subjected d’ as well as c values to a mixed repeated-measures ANOVA design with the between-participants factor *AGE* (younger/older) as well as the within-participants factors *STIMULUS* (unimodal/congruent/incongruent) and *TACS* (sham/beta/gamma). In addition, bias was tested against zero to check whether there would be any statistically significant response tendency at all.

### Correlation analysis

Predictors of cognitive status were evaluated by linear correlation of the DemTect score with age, sensory acuity (see [4] below), grand averages of RTs, d’ and criterion c as well as the effect of multisensory congruence enhancement (MCE; see [5])^14, 34^ on d’ and the effect of target presentation on RTs (working memory enhancement, WME hereafter; see [6]). If both variables did not violate normality, we estimated Pearson’s correlation coefficient, otherwise we used Spearman’s rank correlation. Additionally, we computed all possible bivariate correlations between predictor variables and finally computed partial correlations of each predictor to DemTect controlling for any of the other predictors.

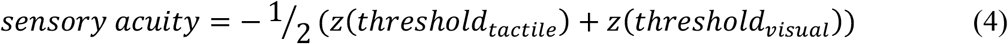

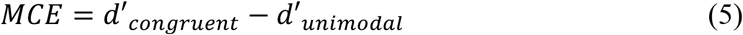

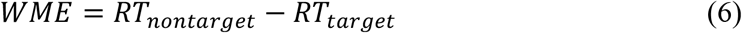

In (6), RT refers to the probability density difference in individual CDF at the RT latency of maximal TARGET effect.

### tACS-related adverse events

The questionnaire tested the perceived maximum intensity as well as the time course of the following sensations: skin sensations (itching, warmth, stinging, pulsating), phosphenes, fatigue and pain (ranked as either “absent”/0, “light”/1, “moderate”/2, “pronounced”/3 or “strong”/4). The time course was evaluated as “beginning”, “end”, “always”. Differences in sensation intensity between age groups as well as between experimental conditions were evaluated with Wilcoxon signed rank tests for zero median. The effectiveness of blinding was assessed by comparing the time course of sensation between sham and verum (beta and gamma) stimulation. To that end, we computed binary compound scores across all sensation qualities that reflected whether participants rather perceived stimulation only in the beginning (0) or at any or all later time points (1). These binary scores were evaluated for significant differences between sham and verum using McNemar tests.

## Acknowledgements

This work was funded by the German Research Foundation (DFG) and the Natural Science Foundation of China (NSFC) in projects SFB TRR169/A3/B1/B4 and by the German Research Foundation (DFG) in projects SFB 936/A3/C1 and SPP 1665/EN 533/13-1.

## Supplementary information

**Figure S1.**
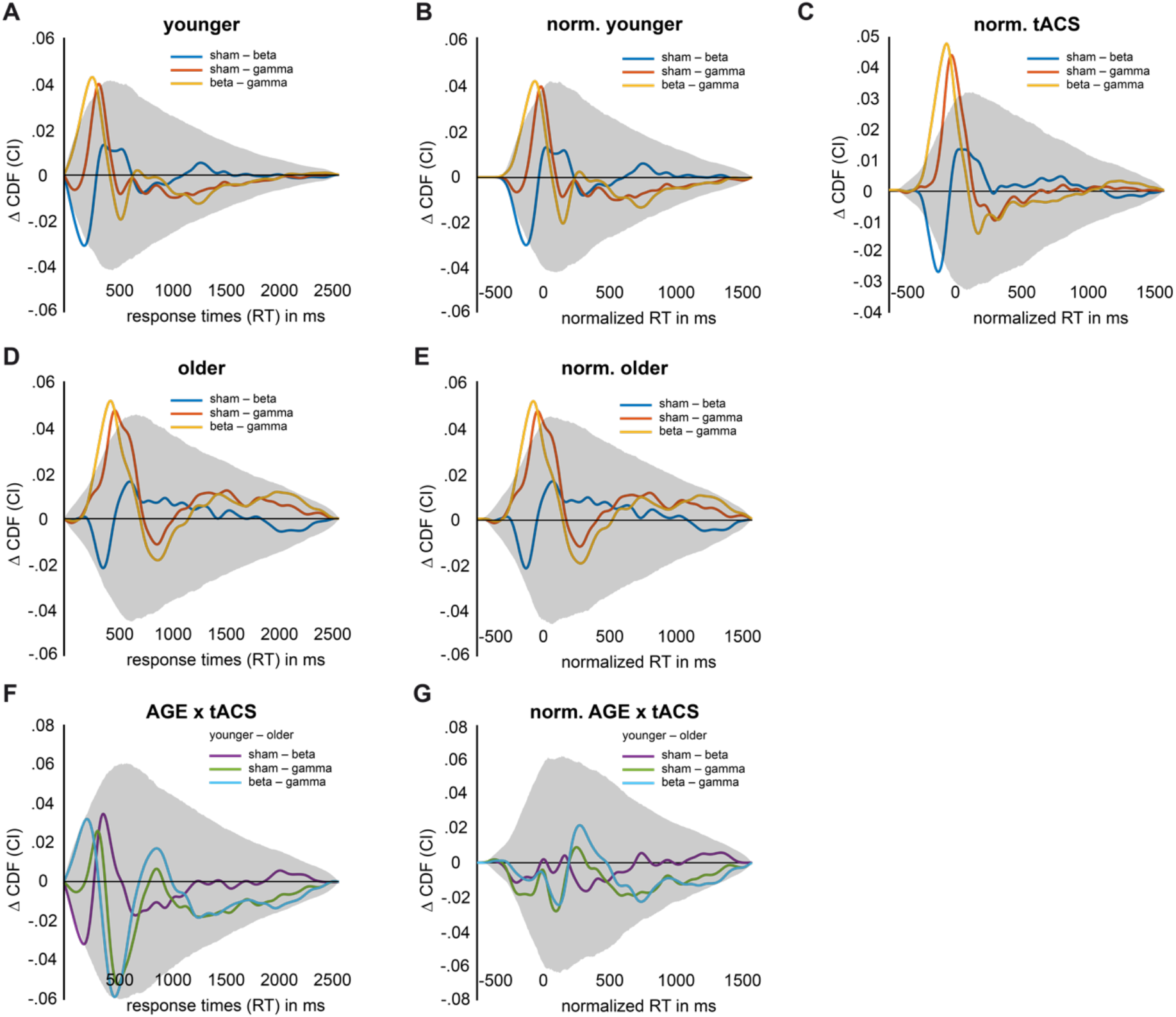
No meaningful interaction between tACS and AGE. Gray shaded area depicts corrected confidence interval. **(A)** Simple effects of tACS in younger participants. **(B)** Simple effects of tACS in younger participants for normalized cumulative distribution functions (CDFs). **(C)** Main effect of tACS for normalized CDFs. **(D)** Simple effects of tACS in older participants. **(E)** Simple effects of tACS in older participants for normalized CDFs. **(F)** Interaction effect between AGE and tACS. **(G)** Interaction effect between AGE and tACS for normalized CDFs.

**Figure S2.**
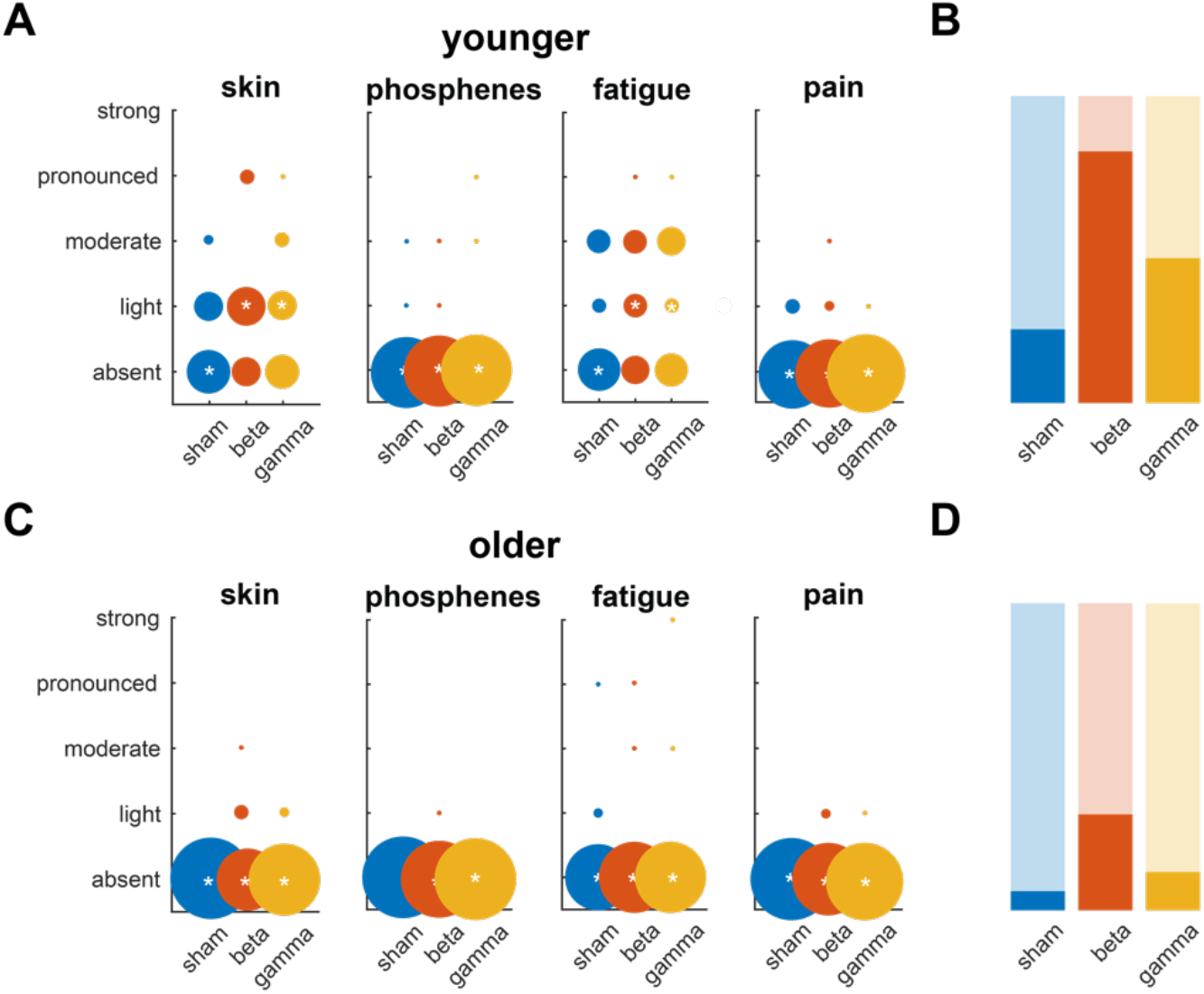
Questionnaire data of tACS side effects shown separately for skin sensations, phosphenes, fatigue and pain. **(A, C)** Younger (A) and older (C) participants rated maximum intensity of a given sensation for each stimulation block (sham, beta and gamma) from absent to strong. The size of circles indicates count of rating and median rating is marked with an asterisk. (**B, D)** Younger (B) and older (D) participants rated whether sensations were present only in the beginning (darker shade) or at any / all later timepoints in the blocks (lighter shade).

